# Exploiting the sequence diversity of TALE-like repeats to vary the strength of dTALE-promoter interactions

**DOI:** 10.1101/068908

**Authors:** Orlando de Lange, Niklas Schandry, Markus Wunderlich, Kenneth Wayne Berendzen, Thomas Lahaye

## Introduction

TALE (Transcriptional activator like effector) repeat arrays are a popular form of programmable DNA binding domain. Reprogrammed TALE transcription factors are referred to as dTALEs (designer-TALEs), and are used within the field of synthetic biology^1^ as well as fundamental research^2^.

Naturally TALEs are pathogenicity factors secreted by *Xanthomonas spp.* into host plant cells to bind specific promoters and manipulate host transcription^3^. Sequence specific DNA binding is conferred by the TALE repeat array. Natural TALE repeat arrays are formed of 10-30 repeats^4^, each of which pairs with one DNA base. Collectively the repeats of a TALE array form a right-handed superhelix, enfolding the DNA, allowing repeats to contact their target bases ^5,6^. Although each repeat is 33-35 amino acids long, polymorphisms are mostly restricted to the residues occupying positions 12 and 13, termed the Repeat Variable Di-residue (RVD). RVDs are oriented in close proximity with target bases^5,6^ and they define the base preference of each repeat^7,8^. The sequence of all RVDs from the N to the C-terminal end of the repeat-array, known as the RVD composition, determines the 5’-3’ DNA sequence specificity. The flanking residues of each repeat are termed non-RVDs and are generally seen as fixed scaffolds to house RVDs. Natural *TALE* genes seem to evolve rapidly with respect to the number and RVD composition of the repeats they encode, forming TALEs with new target preferences^9^. Non-RVDs are contrastingly conserved, a factor that may speed the evolution of *TALEs* through repeat recombinations^10^.

dTALEs, like natural TALEs, mostly differ in repeat number and RVD composition, with non-RVDs held constant. In most currently available dTALE assembly kits, repeat arrays are assembled from the four most common naturally-occurring RVDs. In these toolkits RVD NI (Asn at repeat position 12, Ile at position 13) is used to target A, RVD HD for base C, NG for T and NN for G^4^. This keeps dTALE design simple but does not exploit the full potential of the TALE repeat array for DNA binding. Extensive studies on uncommon or unnatural RVDs have been undertaken and revealed new base specificities^11,12^. In fact the base preferences of all 400 possible amino acid pairs have been assayed in dTALE repeats^13^.

In this study we explore the potential for non-RVD polymorphisms as additional parameters in dTALE design. TALE structural data show that non-RVDs do not contact DNA bases^5,6^. They do, however, determine the superhelical shape of the repeat array through interactions between non-RVDs of each repeat and those of neighboring repeats^5,6^. In addition, non-RVDs can make contacts with the DNA backbone, contributing to the overall affinity of the dTALE-DNA interaction^5,6^.

The power of non-RVDs to modify DNA binding parameters other than base-specificity has already been demonstrated in a few cases. A recent study showed that repeat polymorphisms at two non-RVD positions can increase the binding affinity between a dTALE repeat array and its target DNA^14^. Another study showed that extended TALE repeats, with additional non-RVD amino acids, can be used as a tool to provide optional target base skipping^15^. Both of those studies drew their inspiration from natural repeat sequence variants found among TALEs of *Xanthomonas spp.*. The advantage of sticking to natural variation is that it limits the non-RVD sequence space for testing to those that have been pre-screened by natural selection. Unfortunately, as mentioned, there is little non-RVD variation among the repeats of *Xanthomonas* TALEs. However, TALE-like proteins are produced by other bacteria, including members of the *Ralstonia solanacearum* species complex, whose TALE-likes show far greater non-RVD polymorphism than those of TALEs^10^. TALE-likes are also found in *Burkholderia rhizoxinica*^16,17^ and two unknown marine bacteria^18^. Non-RVD polymorphisms abound in TALE-like repeats at almost every position (Figure S1), yet in all studied cases RVDs confer the same specificities in these TALE-like repeats as they do in *Xanthomonas* TALE repeats. Insights from studies on TALE repeat array structures and these observations from comparative studies of TALE-likes inspired us to use natural non-RVD polymorphisms to vary repeat array binding strength while keeping target sequences constant.

We used the natural pool of sequence-diversity in TALE and TALE-like repeats to assemble sequence diverse repeat arrays, termed variable sequence TALEs (VarSeTALEs). The VarSeTALEs we created have conserved RVD compositions but differ substantially at non-RVD positions. For this work we used sequences from previously characterized TALE-like proteins from *Ralstonia solanacearum*^10,19^ and *Burkholderia rhizoxinica*^17^ strains. We also drew on the full diversity of *Xanthomonas* TALEs, which is generally not used (most dTALEs previously published are derived from two TALEs: AvrBs3^20^ and Hax3^12^).

Our design goal for this study was to generate dTALEs that target the same DNA sequence but do so with a range of binding affinities. We indeed observed that VarSeTALEs mediate a range of promoter activation or repression levels in reporter assays. This is, to the best of our knowledge, the first report on the use of natural TALE-like sequence diversity to tune activities of dTALE repeat arrays while keeping RVD composition constant.

## Methods

### VarSeTALE design

Intra-repeat VarSeTALEs were designed by randomly selecting sets of sequences from a set of unique TALE, RipTAL and Bat sub-repeat modules (Table S1), corresponding to secondary structural elements based on alignments to solved TALE and TALE-like repeat array structures^5,6,21^. Each Intra-repeat VarSeTALE contains a block of 3 or 4 such randomly-assembled repeats, replacing an equal number of AvrBs3 repeats at positions 1-4, 5-7 or 7-B. See Figure S2 for sequences and further details.

Inter-repeat VarSeTALEs were designed by randomly selecting from a set of unique TALE, RipTAL and Bat whole repeat sequences (Figure S2). Each Inter-repeat VarSeTALE contains a block of 5 or 10 such repeats, replacing an equal number of AvrBs3 repeats at positions 1-5, 6-10 or 1-10. Inter-repeat VarSeTALEs 5 and 6 are the combinations of 1 and 3, and 2 and 4 respectively. See Figure S2 for sequences and further details.

All *VarSeTALE* repeat blocks were synthesized (Genscript) with *Xanthomonas euvesicatoria codon* usage and flanked by BpiI restriction sites to facilitate assembly into *dTALE*s as described previously^19^.

The RVD compositions of all dTALEs and VarSeTALEs in this study were chosen to recognize the sequence of the natural AvrBs3 target box from *Capsicum annuum* gene *Bs3*. Multiple RVDs are known to recognize Adenine bases^13^, and for this reason the RVD composition differs slightly between VarSeTALEs. Specifically the RVDs of repeats 1 and 3 differ between inter-and intra-repeat VarSeTALEs and thus separate reference dTALEs are provided for each.

### Molecular cloning

For the repressor assays displayed in Figure 2 *VarSeTALE* repeat arrays were cloned into a derivative of E. coli expression vector *pBT102* bearing truncated AvrBs3 N- and C-terminal domains, via Golden Gate cloning as described previously^18^. The promoter sequence of the cognate reporter (Figure S4) was introduced into *pSMB6* via PCR as previously described^18^.

For protoplast activation assays *VarSeTALE* repeat arrays were cloned into a *pENTR/D-TOPO* derivative containing an *avrBs3* CDS lacking repeats with BpiI restriction sites in their place, as described previously^19^. *CDSs* of VarSeTALEs were then moved into T-DNA vector *pGWB605*^22^ via Gateway LR reaction (ThermoFischer Scientific). The resulting gene is a *CaMV35-S* promoter driven 3’ *GFP* fusion. The reporter was the 360bp fragment of the *C. annuum Bs3* promoter cloned into *pENTR-Bs3p-mCherry* (Figure S4).

### *E. coli* repressor assay

The assay was carried out as described previously^18^. Briefly, *TALE* genes and *mCherry* reporter genes, carried on separate plasmids and driven by different constitutive promoters, are co-transformed into *E. coli (TOP10)* cells. Colonies were allowed to grow to saturation on plate for 24 hours and then single colonies were used to inoculate 150µl scale liquid cultures in 96-well clear-bottom plates. Optical density at 600 nm and mCherry fluorescence were measured after 3.5 hours growth using a Tecan Safire2 plate reader and used to calculate a repression value for each construct, comparing in each case to the combination of the reporter with a dTALE lacking any binding site in the reporter.

### Protoplast transfections & flow cytometry

Arabidopsis root cell culture protoplasts were prepared and transfected as described ^10^. 3µg of *35-S:: TALE-GFP* plasmid was co-transfected with 5µg of mCherry reporter plasmid. The reporter gene was downstream of the *Bs3* promoter which exhibits low basal expression in plant cells^3^, contains the binding site of TALE AvrBs3, used as the basis for all dTALEs in this study. The DNA binding domain of the negative control dTALE has no cognate binding site in the *Bs3* promoter. GFP and mCherry fluorescence were measured in a MoFlo XDP (Beckman Coulter) with a separate blue (488nm, elliptical focus) and yellow (561nm, spherical focus) laser for each fluorophore. GFP peak emission was captured by a 534/30 bandpass, mCherry peak emission by a 625/26 bandpass. Viable cells were identified by gating out dead cells by comparing narrow-scatter log-area vs. large-angle scatter log-area. This was followed by elimination of large cell clumps by comparing large-angle scatter log-area to large-angle scatter pulse width. Thereafter each GFP population was identified as cells having more fluorescence emission in the FL1 (534/30) compared to the FL2 (585/29) over that of un-transfected cells. Similar, mCherry expressing cells were identified by comparing FL7 (625/26) to FL6 (580/23). Finally, a gate [GFP or mCherry] was made to capture all transfected cells and exported.

Using log(mCherry) as a response variable, and log(GFP) as a proxy to VarSeTALE expression, the mean log(mCherry intensity) per log(GFP intensity) was estimated for each construct using a linear model restricted to cells expressing [GFP, log(GFP)≥1.8]; at least 1500 cells (mean 3000 cells) per biological replicate were obtained. The mean increase from at least 2 (3 in most cases) biological replicates of log(mCherry)/log(GFP) relative to the negative control with 95% confidence intervals are shown in Figure 3. Analysis was performed using R, and the lm function from package stats was used to construct a linear model. Pairwise differences were calculated using the multcomp package^23^.

### Plant material and *Agrobacterium* leaf infiltrations

Pepper (*Capsicum annuum*) plants of cultivar ECW–30R containing the resistance gene *Bs3* were grown in the greenhouse at 19°C, with 16 h of light and 30% humidity. Vector constructs were introduced into *Agrobacterium tumefaciens* strain GV3101 pMP90 by electroporation and selection on YEB medium^24^ containing the appropriate antibiotics. *Agrobacterium* strains were grown as liquid culture for 24 hours in YEB medium, harvested by centrifugation and resuspended in sterile water at an OD of 0.4 for infiltration. The suspension was injected into the lower side of leaves from six-week-old pepper plants. After 48 hours infiltrated patches were cut out and stored at −80°C for RNA extraction.

### Isolation of RNA and quantitative real-time RT-PCR (qPCR) analysis

RNA was isolated from 50 mg frozen leaf powder with the GeneMATRIX Universal RNA Purification Kit (EURX, Gdansk, Poland). Reverse transcription was performed with 1 μg of the total RNA using the iSCRIPT cDNA Synthesis Kit (Biorad, Hercules, CA). Quantitative PCR reactions were performed using SYBR^®^ Green technology (MESA GREEN qPCR Mastermix, Eurogentec, Germany) on an Bio-Rad CFX384 system (Biorad, Hercules, CA). *Bs3* cDNA was amplified with primers Bs3 RT F7 and Bs3 RT R7, EF1-α cDNA with primers EF1a F2 and EF1a R2, ß-TUB cDNA with the primers ß-TUB F2 and ß-TUB R2. Data were analyzed employing the Bio-Rad CFX Manager 3.1 software with EF1-α or β-TUBULIN as a reference gene.

## Results

### VarSeTALE repeat arrays contain large numbers of non-RVD polymorphisms

In this study we generated variable sequence TALEs (VarSeTALEs), which are dTALE repeat arrays bearing several repeats with sequences drawn from different TALE and TALE-like origins. Specifically we generated VarSeTALEs repeat modules using sequences from *Xanthomonas* TALEs, *Ralstonia* TALE-likes (RipTALs^25^) and *Burkholderia* TALE-likes (BurrH/Bat1 and Bat2^16,17^). Sequences of TALE-likes repeats used as the raw material for design for the repeat arrays generated in this study are displayed in Figure S1.

We explored two alternative approaches for VarSeTALE design. We combined either whole repeats (inter-repeat VarSeTALEs) or repeat subunits (intra-repeat VarSeTALEs). For inter-repeat VarSeTALEs the highly conserved leucine residue at position 29 (Figure S1) was used as the breakpoint between repeats of different origins. Repeat subunits used in our intra-repeat VarSeTALEs correspond to secondary structural elements: short-helix (4-10), RVD loop (11-15), long-helix (16-28) and inter-repeat loop (29-1). Figure 1 illustrates the two design approaches using example sequences.

**Figure 1:**
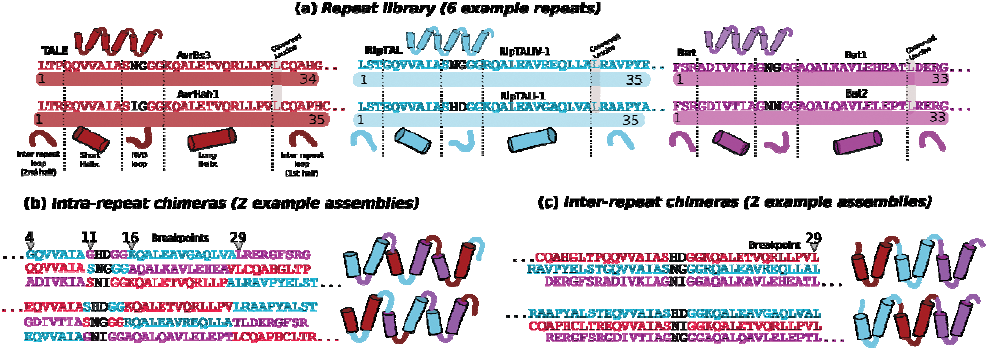
VarSeTALE design. (a) Starting material was an *in-silico* repeat library of non-identical repeat sequences from TALEs and TALE-likes. Sequences were sorted based on bacterial origin (TALEs of *Xanthomonas*, RipTALs of *Ralstonia solanacearum* and Bats of *Burkholderia*). Color coding reflects these groupings throughout this figure. Numbers indicate residue positions within each repeat, as classically defined. Throughout this figure RVD residues are left uncolored, as RVDs are kept constant. To facilitate intra-repeat VarSeTALE design, the known TALE repeat structure^5,6^ was divided up into predicted secondary structural elements. To facilitate inter-repeat VarSeTALE design, we searched for a repeat position, close to a helix-loop transition, that is conserved across all TALE-like repeats within the library; this was leucine 29. Repeat subunits (b) or whole repeats (c) were randomly shuffled to design sequences encoding blocks of sequence diverse repeats. These repeats were synthesized and cloned into otherwise standard dTALE repeat arrays, generating VarSeTALEs for functional testing.

The VarSeTALEs generated in this study are AvrBs3 derivatives, bearing 3-10 sequence diverse repeats in place of AvrBs3 repeats. In the case of intra-repeat VarSeTALEs only 3-4 repeats per array were replaced, whereas 5-10 were replaced to create inter-repeat VarSeTALEs. All repeat arrays generated have an RVD composition identical or equivalent to that of AvrBs3. All intra-repeat VarSeTALEs have exactly the same RVD composition as AvrBs3, while intra-repeat VarSeTALEs use NI RVDs (A-specifying) in two positions where other A-specifying RVDs are found in AvrBs3. Please refer to materials and methods for further details and figure S2 for full amino-acid sequences of all VarSeTALEs generated.

We hypothesized that the non-RVD polymorphisms of VarSeTALEs will result in differing binding strengths on their target DNA boxes. We used three experimental approaches that infer relative binding strengths from differential promoter regulation: An *E. coli* promoter repression assay (Figure 2), an *Arabidopsis* protoplast transactivation assay (Figure 3) and *Agrobacterium* delivery into *Capsicum annuum* (bell pepper) to activate a genomic promoter (Figure 4).

### *Differential promoter repression by VarSeTALEs in* E. coli

The first approach we used to compare activities of VarSeTALEs and reference TALEs was a repression assay in *E. coli*, based on a TALE-repressor system^26 18^. In this assay a TALE binds to a modified *Trc* promoter driving constitutive *mCherry* expression in *E. coli*. dTALE-promoter binding is assumed to impair promoter activity by occluding the RNA polymerase complex. We were previously able to demonstrate that in this assay repression correlates to DNA-binding affinity as measured *in vitro*^18^. VarSeTALE and reporter plasmids were co-transformed into *E. coli* and resulting colonies were used to inoculate separate cultures in wells of a 96-well plate. After 3.5 hours of further growth mCherry expression and cell density (OD 600) were measured in a plate reader. Results are shown in Figure 2.

Our expectation was that VarSeTALEs would mediate a range of reporter activities. No prediction was made as to the activities of individual VarSeTALEs. Instead, we expected that due to the spread of sequence polymorphisms the whole set of VarSeTALEs would capture a range of reporter repression levels. That is indeed what we observed (Figure 2). For both the intra- and inter-repeat VarSeTALEs the range of repression strengths ranged from barely detectable to above the activity of the reference dTALE, as inferred from comparison of sample medians.

**Figure 2:**
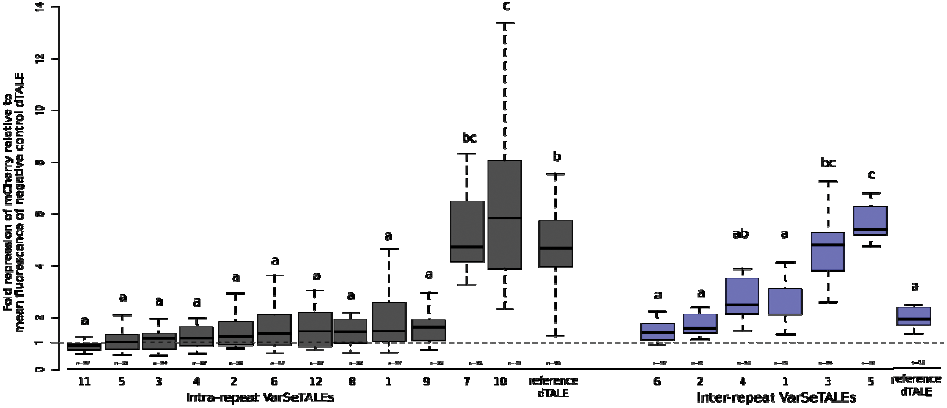
Ability of a set of VarSeTALEs and cognate reference dTALEs to repress a promoter driving an mCherry reporter in *E. coli* cells. 13 intra-repeat (gray) and 6 inter-repeat (blue) VarSeTALEs were assembled based on a pool of diverse TALE repeat sequences. All VarSeTALEs and control dTALEs were tested for their ability to repress transcription from a bacterial promoter containing a cognate binding element, in a promoter driving expression of an mCherry reporter. Boxplots of fold *mCherry* promoter repression (Base10), relative to an unrelated, negative control, dTALE (Supplementary Figure 2C) are shown along with the number of biological replicates tested given underneath in each case. A dotted line at 1 indicates basal reporter activity without repression. Each biological replicate corresponds to a single colony picked into a 96-well assay plate. Reference dTALEs were assembled entirely from AvrBs3-derived repeats^20^. VarseTALEs are ordered within their groups based on increasing repression strength, with numbers (X) below each boxplot giving the identifier of each VarSeTALE (IntraX or InterX in Table S1). Letters above boxplots indicate significance groups derived generalized linear hypothesis testing conducted separately for inter- and intra-repeat VarSeTALEs; samples sharing a letter are not significantly different.

For both designs a range of repression strengths were achieved but the range was smaller for inter-repeat VarSeTALEs. Ten out of thirteen intra-repeat VarSeTALEs (Figure 2, gray) mediated significantly weaker reporter repression than the reference dTALE, though intra-repeat VarSeTALE 10 mediated significantly stronger repression than the reference. The inter-repeat VarSeTALEs displayed a similar relationship to their reference dTALE but with a slightly smaller total range of median fold repression strengths: 4.1 compared to 4.8 for intra-repeat VarSeTALEs.

Since ten of the Intra-repeat VarSeTALEs displayed repression strengths that were not significantly different from one another some were set aside in the next experiments. Intra-repeat VarSeTALEs 1, and 4-10 were chosen to capture the full range of activities measured for the repressor reporter (Figure 2).

### *Differential promoter transactivation by VarSeTALEs in* Arabidopsis *protoplasts*

The *E. coli* repression assay used in figure 2 gives a straightforward read out of stoichiometric promoter repression, which should correlate directly to DNA binding affinity. However, when dTALEs are used in eukaryotes for regulation of synthetic genetic circuits, they are fused to activation or repression domains^27^. In this context the relationship between promoter regulation and DNA binding is less direct. So we next tested the ability of VarSeTALEs to activate a promoter driving a fluorescent reporter in eukaryotic cells (Figure 3). We chose to work in *Arabidopsis* root cell-culture protoplasts to exploit the natural C-terminal domain of AvrBs3, which encodes a strong *in planta* transactivation domain^28^. Each VarSeTALE was GFP-tagged to allow us to monitor VarSeTALE expression and then derive a statistical estimate of the transactivation strength for each VarSeTALE (see Materials and Methods and supplementary material S4 for further details).

**Figure 3:**
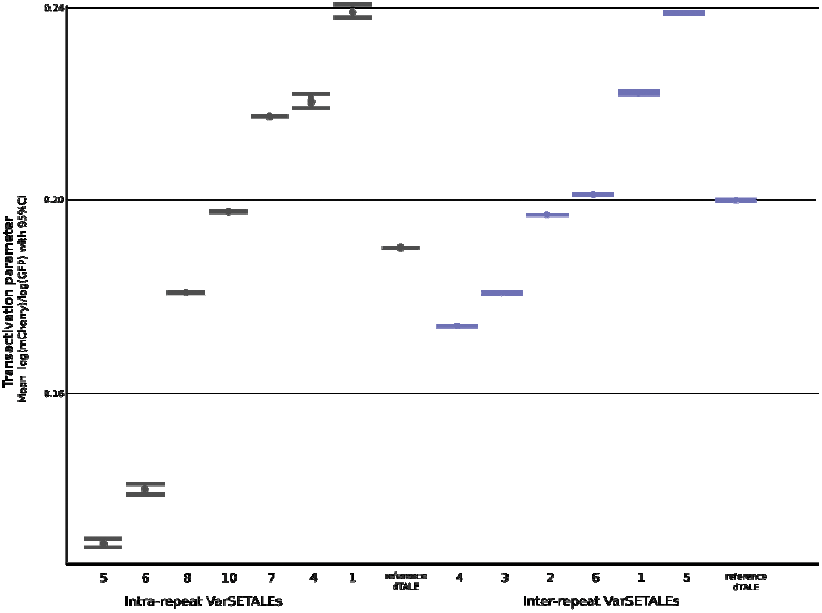
Reporter activation by a set of VarSeTALEs, measured in Arabidopsis protoplasts. 7 intra- (gray) and 6 inter- (blue) repeat VarSeTALEs, tagged with GFP, were transfected into *A. thaliana* protoplasts alongside an *mCherry* reporter gene containing a cognate binding site in its promoter (*pBs3*). Fluorescence levels were measured using a MoFlo XDP flow cytometer. From these data VarSeTALE promoter activation parameters were estimated using a linear model, as described in materials and methods. The estimated parameter values as an increase relative to dTBat1 (the negative control), are displayed here along with 95% confidence intervals. VarSeTALEs are ordered within their groups based on increasing activation strength (slope mCherry/GFP), with numbers giving the identifier of each VarSeTALE (IntraX or InterX in Table S1). Statistical analysis was conducted separately for inter- and intra-repeat VarSeTALEs; samples whose mean is not within the confidence interval of another are significantly different (alpha=0.05).

As for the repressor assay (Figure 2) we found that VarSeTALEs mediated a range of transactivation strengths (Figure 3). This time the difference between intra- and inter-repeat VarSeTALEs was more pronounced. The seven intra-repeat VarSeTALEs we assayed spanned a range of transactivation parameters with the same maximum and a lower minimum than the inter-repeat VarSeTALEs.

Interestingly, the relative performances of specific VarSeTALEs often differed in the transactivation assay and repressor assay. Intra-repeat VarSeTALE 5 and inter-repeat VarSeTALE 5 do both occupy the same relative positions, as the worst and best performers, respectively, in both assays. For all other constructs there is no obvious connection between repression performance and transactivation performance. This is perhaps not surprising when one considers the conceptual difference between an assay of stoichiometric repression and one of promoter transactivation. A strong VarSeTALE-DNA interaction may lead to strong stoichiometric repression^18^ (Figure 2). By contrast, promoter activation involves recruitment of the transcriptional machinery and unwinding of the double helix coupled to strand-disassociation downstream to allow transcription. In such a scenario a high affinity, particularly a low K_off_ may be disadvantageous. A study that derived DNA binding affinities as well as fold-activations for a set of 20 dTALEs, differing in RVD composition, found an overall positive correlation between DNA binding affinity and promoter activation, but this correlation disappeared for the highest affinity TALE-DNA pairings^29^. Thus some of the observed discrepancies are likely a consequence of the differences between stoichiometric repression and promoter transactivation.

### *Differential genomic promoter activation by VarSeTALEs in* Capsicum annuum *leaf cells*

The key specification we were hoping to achieve from our designs is that VarSeTALEs with the same RVD composition can bind and regulate a promoter to a range of levels, and in this they achieved their aim. We therefore next tested whether this property was preserved in the activation of a chromosomally embedded gene, a common application of dTALEs^2,30^. The *Bs3* gene of bell pepper (*Capsicum annuum* ECW30-R) contains a target site for AvrBs3 in its promoter^3^. We introduced CaMV35-S promoter driven *VarSeTALE* genes into bell pepper leaves via *Agrobacterium tumefaciens* transient transformation and quantified *Bs3* transcript levels via qPCR, which provides a proxy for promoter activation levels (Figure 4).

**Figure 4:**
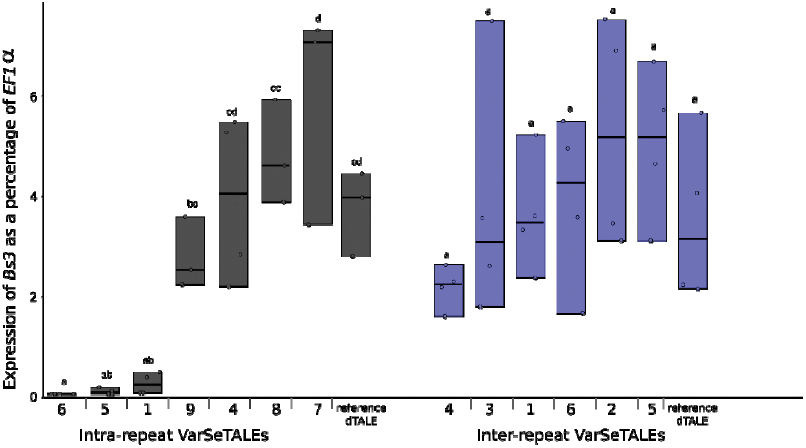
A range of gene expression levels achieved for a *Capsicum annuum* endogene by using a set of VarSeTALEs. Leaves of *C. annuum* plants containing the *Bs3* gene, a natural target of TALE AvrBs3 and therefore of the VarSeTALEs in this study, were infiltrated with *A. tumefaciens* strains to deliver VarSeTALEs and reference dTALEs. Expression of the *Bs3* gene was quantified with qPCR and compared to housekeeping gene EF1a (Elongation factor 1 alpha). Circles represent individual replicates, which are connected into a datablock with a thick black line at the sample median. Numbers (X) below each datablock giving the identifier of each VarSeTALE (IntraX or InterX in Table S1). Letters above indicate significance groups based on pairwise t-tests; samples sharing a letter are not significantly different. Absolute values for the intra- and inter- repeat VarSeTALEs are likely influenced by the performance of the assays on separate days on separate plants.

We expected to see a range of activation levels of the usually transcriptionally silent *Bs3* gene. This is indeed what we observed (Figure 4), with VarSeTALEs of both design types. However, only in the case of the intra-repeat VarSeTALEs were the differences we observed statistically significant. This is likely to be to a great extent a reflection of the high variability between replicates, arising from the variation in *Agrobacterium* infection and DNA delivery between leaf samples. However, it is consistent with previous assays that the intra-repeat VarSeTALEs showed a greater range of promoter activation strengths.

Again the relative performances of VarSeTALEs measured in this assay do not correspond well to the results of the transactivation reporter assay (Figure 3). In this case both assays provide a measure of promoter transactivation in plant cells. Interestingly the weakest activators in both transactivation assays (intra-repeat VarSeTALEs 5 and 6, and inter-repeat VarSeTALE 4) are also among the weaker repressors (Figure 2). This suggests that those VarSeTALEs are poor DNA binders leading to consistently weak promoter regulation. The dataset in this study is not extensive enough to allow detailed analysis of the effects of specific sets of non-RVD polymorphisms even though our overall results do indicate that VarSeTALE repeat arrays, containing high numbers of non-RVD polymorphisms, bind DNA and regulate the same promoters to different levels.

## Discussion

### VarSeTALEs harness non-RVD polymorphism to tune promoter regulation

Our goal was to harness natural non-RVD polymorphisms as a means to vary TALE-DNA binding affinity without changing base preference. We created sequence diverse dTALE repeat arrays, termed VarSeTALEs, by drawing on natural TALE and TALE-like repeat diversity. We combined either whole repeats (inter-repeat VarSeTALEs) or repeat subunits (intra-repeat VarSeTALEs) to create the sets of sequence diverse repeats used in the study. Using three different experimental approaches we demonstrated that sets of six to twelve intra or inter-repeat VarSeTALEs can regulate the same target promoter to different levels (Figures 2-4). The observed differences in promoter activity are consistent with a range of VarSeTALE-DNA binding affinities based on previous work with the *E. coli* TALE-repressor assay directly comparing repression with DNA binding affinity ^18^. Our data thus support the VarSeTALE approach as a general method to vary TALE-DNA affinity while keeping RVD composition and thus DNA target sequence constant.

### Comparing VarSeTALE design approaches: intra vs inter-repeat

We used two different approaches to design VarSeTALEs (Figure 1) but in each case the goal was the same: to vary binding strength whilst retaining target sequence recognition. Sets of both intra- and inter- repeat VarSeTALEs mediated a range of repression (Figure 2) or activation strengths (Figures 3 and 4). However, the intra-repeat constructs consistently outperformed the inter-repeat VarSeTALEs because they covered both a greater absolute range and mediated effect strengths both above and below that of their reference dTALE in each different assay (Figures 2-4; gray). In contrast inter-repeat VarSeTALEs tended to more closely match the performance of their cognate reference dTALE (Figures 2-4; blue). The intra-repeat VarSeTALE design approach seems to have better achieved the goal of varying repeat array binding strength.

The larger observed effect range of sets of intra-repeat VarSeTALEs compared to inter-repeat VarSeTALEs may stem from the greater number of repeat sequence origins they represent. Each intra-repeat VarSeTALE repeat was assembled from four different subunits: short-helix, long-helix, RVD loop, and inter-repeat loop (Figure 1). Each of those subunits is derived from a different TALE or TALE-like. The non-RVDs of a given TALE or TALE-like repeat array have evolved together, and it seems reasonable to assume that bringing together sequences from phylogenetically distant repeat arrays would disrupt natural intra-molecular interactions. We can assume that most novel combinations of non-RVD polymorphisms will disrupt interactions that normally hold together the TALE-like repeat structure leading to poorer DNA binding. This assumption is supported by previous work showing that rearrangements of the highly polymorphic repeats of Bat1 often impaired repeat array function^17^. Indeed, out of our initial set of 13 intra-repeat VarSeTALEs, most were very poor repressors, not significantly different from the negative control dTALE (Figure 2; dotted line). However, a recent study has also demonstrated that non-RVD polymorphisms that disrupt inter-repeat interactions can increase the structural flexibility of a dTALE repeat array superhelix, enhancing DNA binding^14^. Intra-repeat VarSeTALE repeat arrays contain a greater number of novel non-RVD residue pairings than inter-repeat VarSeTALEs which may explain the diversity of promoter regulation strengths we observed for these constructs.

### Applications for VarSeTALEs: Controlling synthetic gene circuits, reverse genetics and transgene stability

The creation of synthetic genetic circuits is a central practice of synthetic biology^31^. Promoters are used as key regulation points within synthetic genetic circuits, tuning circuit flux through downstream gene expression. They also serve as integration points for inputs from other genes encoding transcription factors. By now numerous studies have explored the potential for dTALE-promoter interactions to regulate synthetic genetic promoters, creating analog^26^ or digital^1^ control of gene expression as well as Boolean logic gates^32^. Unsurprisingly, therefore, libraries of TALE-promoter pairs with different binding affinities have been characterized to serve as reusable modules in synthetic genetic circuit design^33,34^. We believe that VarSeTALEs make a useful addition to those existing dTALE tools, filling a slightly different role. VarSeTALEs would be useful in cases where promoter sequence cannot be altered but additional tuning is still desirable. VarSeTALEs could be added to existing synthetic genetic circuits without requiring any redesign of constituent promoters.

Reverse genetics could be another application of VarSeTALEs. In this approach the expression of a gene of unknown function is modified to observe effects on phenotype and therefore gain insights into gene-function. Sets of VarSeTALEs could be built to target the same native promoter with different activation or repression strengths. If a permissive promoter position has been identified a set of VarSeTALEs could be transformed into the organism of interest, rapidly producing at a set of transgenic lines differing in expression of the native gene of interest. This approach would be applicable for activator or repressor dTALEs, both of which have already been used in a range of host organisms^11,33,35^.

An additional benefit of VarSeTALEs is that the DNA sequences encoding their repeats are more diverse. The runs of DNA repeats that encode conventional TALE repeat arrays are problematic for PCR based manipulation^36^ and are susceptible to recombinatorial sequence deletion in some systems^10,37^. In the later case, the problem of recombination can be alleviated by lowering repeat sequence similarity^38^ through codon redundancy, but the added diversity that comes from amino acid level polymorphism provides an alternative solution. Where *dTALE* genes are intended to remain stably as transgenes over multiple generations VarSeTALEs may serve better than conventional dTALEs.

### Future improvements to VarSeTALE design

We envision that the VarSeTALEs assembled in this study as an initial proof-of-concept and encourage the development of better tools to search within the total VarSeTALE design space. The VarSeTALEs in this study only capture a small subset of TALE and TALE-like repeat diversity. Especially since the recent characterization of TALE-like DNA binding proteins from marine bacteria^18^ further expands the sequence pool of TALE-like repeats. The number of possible combinations of TALE and TALE-like repeats and repeat subunits is huge and random searches are a very slow method to arrive at those with desired DNA binding properties. High-throughput assembly and screening could allow selection of promising candidates out of a VarSeTALE repeat library. Alternatively, further study of non-RVDs could be used to build rational design rules, helping users to select promising combinations. For example a recent study used a mix of *in vitro* binding assays and molecular dynamics simulations to understand the functional impact of certain non-RVD polymorphisms at two positions within dTALE arrays^14^. Further studies on the structural and functional impacts of non-RVD polymorphisms could allow improved VarSeTALE design.

A core assumption of our design approach was that non-RVDs alter overall DNA binding affinity but do not change the target base preference of TALE repeats. This is based on previous work on TALE-like repeat arrays which the same RVD-target base associations despite considerable non-RVD polymorphism^17–19^. Yet high-throughput screens have shown that the base preference of dTALE repeats is often slightly altered by neighboring repeats^13^. It therefore seems likely that the alterations to intra- and inter-repeat molecular interactions inherent to VarSeTALE design will have a range of subtle effects on base preference. A range of experimental approaches have been developed to screen base preference of dTALE repeat arrays using pools of random oligonucleotides as binding targets^29,39,40^. These methods could be applied to VarSeTALEs to provide more information on base preference. This would be important to accurately predict off-targets in a genomic context.

We encourage further work to explore the VarSeTALE design concept while equally inviting interested parties to use the exact sequences in this study (provided in Figure S2) as chassis for creating novel sets of VarSeTALEs by simply replacing RVDs used here with those matching a DNA target of interest. We would stress however, that upon generating VarSeTALEs with a new RVD composition that their relative performances should be tested in the system of interest, since, as we have shown, relative activities of some VarSeTALEs differed considerably in the different assay systems we used in this study. However, what we anticipate is that using a set of VarSeTALEs, either those presented here or independently derived, will capture a range of promoter regulation levels without the requirement for any rational engineering.

## SI

Table S1: Unique sub-repeat modules used for intra-repeat VarSeTALE assembly

Table S2: Raw data and accompanying calculations used in the preparation of Figure 2 (*E. coli* repressor assay).

Table S3: Raw data and accompanying calculations used in the preparation of Figure 3 (*Arabidopsis* protoplast transactivation assay).

Table S4: Raw data and accompanying calculations used in the preparation of Figure 4 (*Capsicum annuum* leaf tissue transactivation assay).

Table S5: Primer sequences used for qPCR experiments

Figure S1: Alignment of natural TALE repeat variation used as the basis for VarSeTALE design (PDF)

Figure S2: Sequences of VarSeTALEs and reference dTALEs. (docx Word document)

Figure S3: R Markdown for statistical analysis for figure 2. (PDF)

Figure S4: R Markdown of analysis for figure 3. (PDF)

Figure S5: Sequences of reporter plasmids (pSMB6 EBE AvrBs3 and pENTR-Bs3p-mCherry) (docx Word document)

## Acknowledgements

N. S. was supported by the “Institutional strategy of the University of Tübingen” (DFG ZUK63).

